# Maternal glucocorticoids and behavior shape offspring developmental trade-offs in wild baboons

**DOI:** 10.1101/2025.11.18.689046

**Authors:** Sam K. Patterson, Katie Hinde, Angela Bowen Bond, Benjamin C. Trumble, Amy Lu, Shirley C. Strum, Joan B. Silk

## Abstract

Mammalian mothers provide behavioral and physiological signals that offspring use to calibrate development in relation to maternal resources and environmental cues. Infants respond selectively as they prioritize certain developmental systems over others, creating developmental tradeoffs between competing biological systems. Here, we investigate the influence of maternal capital (“investment capacity”) on the growth and development of their infants in wild olive baboons (*Papio anubis*) from Laikipia, Kenya. We posit that maternal capital is influenced by a mother’s own early life experiences (e.g., drought, maternal loss) and her current life experiences (e.g., dominance rank, food availability), and is signaled to offspring via maternal effort (i.e., nursing and carrying time) and glucocorticoids. We used behavioral data on 40 infants (43% female) in the first year of life to quantify maternal effort, infant play, and infant independence (i.e., frequency of infant departures from mother). We matched these behavioral data with maternal fecal glucocorticoid measures from lactating mothers, and infant growth measures assessed via photogrammetry. Signals of low maternal capital predicted lower rates of infant play, less behavioral independence, and slower growth. There was a negative relationship between the rate of social contact play and growth rate, indicating a developmental tradeoff. Males were more sensitive than females to some of the maternal signals measured in our study. These results add to a growing body of evidence demonstrating that maternal behavioral and physiological signals shape infant development.

## Introduction

For mammalian infants, the mother constitutes the adaptively relevant environment organizing developmental trajectories of offspring. The maternal environment, including the mother’s social, nutritional, and psychobiological status along with the care, nutrition, and signals she provides, influences offspring growth, immune function, cognitive development, social development, and reproductive strategies in a variety of taxa (Anzà et al., 2023, 2025; Baniel et al., 2022; Blomquist, 2013; Catalani et al., 2011; Groothuis & Schwabl, 2008; Guenther et al., 2014; Langley-Evans, 2007; Machado, 2013; Malalaharivony et al., 2021; Mateo, 2014; Moore et al., 2019; Most & Strum, 2020; Mousseau & Fox, 1998; Petrullo et al., 2022; I. A. Schneider-Crease et al., 2022). These effects are expected, given the long dependency period during which infants are exposed to maternal inputs. This dependency also allows the mother to act as a buffer between offspring and the external environment. For example, during famines, pregnant women experience up to a 50% decline in caloric availability, but birth weight of offspring is only reduced by about 10% (Stein et al., 2004; Z. Stein et al., 1975; Wells, 2019).

Maternal signaling and investment are expected to vary as a function of mother’s cumulative capital, or the sum of all the maternal characteristics that underlie investment in offspring (Wells, 2019, 2025). Maternal characteristics like high dominance rank, advantageous early life experiences, prime adult age, and social support contribute to greater maternal capital. Information about maternal capital can be conveyed to offspring via physiological signals that are transmitted across the placenta or through mother’s milk and via behavioral signals (de Weerth et al., 2023; Hinde & Milligan, 2011; Lu et al., 2019; Wells, 2025). For example, maternal glucocorticoids (GCs), which reflect energy mobilization, stress, fertility, and health (Palme, 2019; Sapolsky et al., 2000), are passed to offspring through milk and are hypothesized to signal information about the degree and schedule of maternal investment (Allen-Blevins et al., 2015; Hinde & Milligan, 2011; Lu et al., 2019; Meaney et al., 2007; Pácha, 2000; Wells, 2025). Lower parity rhesus macaques and those with smaller body sizes produce milk with higher GC concentrations, potentially reflecting fewer resources available for offspring (Hinde et al., 2015). Care behaviors like nursing and carrying can also act as maternal behavioral signals. Paradoxically, high levels of maternal effort (i.e., higher nursing and carrying levels) may signal low levels of maternal capital (Altmann & Samuels, 1992). For example, baboon mothers who are low ranking and experienced more early life adversity – i.e., characteristics leading to lower capital – carry and nurse their offspring more than higher ranking females and females who experience less early life adversity (Patterson et al., 2021). Low capital mothers might produce a lower quantity and quality of milk thus resulting in greater maternal effort.

Maternal signals allow offspring to calibrate development based on local conditions and are expected to prioritize certain developmental systems over others, orchestrating developmental tradeoffs. Because of the close connection between mothers and infants during gestation and lactation, mammalian early life development is thought to be primarily in the control of the mother (Wells, 2010, 2019). However, within the limits set by maternal investment, offspring can adaptively allocate energetic resources across different dimensions of development, such as behavior, cognition, immunology, and growth (Allen-Blevins et al., 2015; Lu et al., 2019; Wells, 2019). Offspring born to mothers that provide lower levels of investment might need to cut costs across multiple dimensions of development in order to survive, or they might prioritize some dimensions over others (Allen-Blevins et al., 2015; Lu et al., 2019). For example, in Assamese macaques (*M. assamensis*) elevated prenatal glucocorticoids (GCs) are associated with accelerated postnatal growth but lower rates of play, slower motor skill acquisition, and reduced immune function (Berghänel et al., 2015, 2016). In rhesus macaques, elevated milk GCs are associated with accelerated growth, but greater nervousness, suggesting that higher milk GCs might instruct offspring to prioritize growth over other dimensions of development (Hinde et al., 2015). In both cases, maternal GCs might provide a signal to offspring and shape offspring tradeoffs between growth and development.

As a key life history process, somatic growth represents a dimension of development with fitness consequences (Stearns, 1992). Despite the advantages of accelerated growth and large body size, growth rates vary and are often submaximal (Dmitriew, 2011). Maternal environments influence offspring growth rates (Bernardo, 1996; Mousseau & Fox, 1998; Samuni et al., 2020). In macaques and baboons, greater maternal investment in the form of nutrient transfer is positively associated with offspring growth, and high ranking, prime-aged mothers produce offspring with faster growth rates than other mothers (Altmann & Alberts, 2005; Hinde et al., 2015; Johnson, 2003). Maternal signals like GCs may play a role in guiding offspring growth trajectories. Findings on the relationship between maternal glucocorticoids and offspring growth are mixed, but a meta-analysis indicates that exposure to elevated maternal GCs during early gestation tends to result in accelerated growth, and elevated maternal GCs during late gestation tend to result in slower growth rates (Berghänel et al., 2017). There are very few studies on the effects of milk GCs on infant growth, but milk GCs are negatively associated with infant growth in humans (Hahn-Holbrook et al., 2016). In rhesus macaques, higher milk GCs during early lactation predict slower growth, higher milk GCs at peak lactation predict faster growth (Hinde et al., 2015), and increases in milk GCs from peak to late lactation predict faster growth among daughters, but not sons (Petrullo et al., 2019). These data illustrate the complexities of maternal signals and offspring responses.

Play is a dimension of development that has often been ignored in studies of life history tradeoffs. Social play is a complex activity, involving physical coordination among individuals and is the only activity that simulates the responses and reaction time of real fighting (Graham & Burghardt, 2010). Play is assumed to be widespread in nature because it serves important adaptive functions, such as motor skill development, training for unexpected events, and social benefits that outweigh its energetic costs and risks of injury (Graham & Burghardt, 2010; Palagi, 2018). Rates of play behavior are positively associated with acquisition of motor skills (Belding’s ground squirrels (*Spermophilus beldingi)*:(Nunes et al., 2004); Assamese macaques:(Berghänel et al., 2015)), age at dispersal (Nunes et al., 2004), reproductive success (Nunes et al., 2004), and survival (horses (*Equus caballus)*:(Cameron et al., 2008)). Further, the timing of solo, object, and social play in captive vervets (*Cercopithecus aethiops*) coincides with the timing of development in different parts of the neocortex, suggesting that play behaviors might have permanent effects on the developing brain and adult abilities (Fairbanks, 2000). It is possible that infant play may be constrained by the extent of available energy provided by their mothers. For example, reduced maternal investment is associated with less play in several primate species (Japanese macaque (*M. fuscata)*:(French, 1981); Rhesus macaque (*M. mulatta)*:(Mccormack et al., 2006)), but the opposite pattern has also been reported where reduced investment is associated with more play (domestic cat (*Felis cactus)*:(Bateson et al., 1981, 1990); vervets:(Fairbanks & Hinde, 2013)), perhaps to enable infants to reap the benefits of play before their predicted earlier weaning. Further studies exploring the links between various components of the maternal environment, such as maternal signals, and how offspring prioritize play relative to other developmental systems are needed to contribute to our understanding of these patterns.

Here, we investigate how characteristics and signals associated with maternal capital influence infant development and growth in a wild population of olive baboons (*Papio anubis*) in Laikipia, Kenya. We hypothesize that a mother’s capacity to invest in her offspring will be influenced by her own early experiences in life (e.g. her mother’s rank, her mother’s parity, and environmental conditions at the time of her birth) and her current life circumstances (e.g. current rank, parity, and resource availability). Previous research in this population demonstrated that mothers’ early and current experiences predicted maternal behavioral and physiological signals: more early life adversity, lower rank, and more current challenges like human encounters were associated with greater maternal effort and higher maternal glucocorticoids, whereas greater herbaceous biomass was associated with more maternal effort but lower maternal glucocorticoids (Patterson et al., 2021). We build on this work by assessing the effects of maternal signals (behavior and physiology) and maternal characteristics on offspring outcomes. We examine several predictions derived from this basic hypothesis.

1a. Signals of low maternal capital negatively affect all dimensions of development. Thus, higher levels of maternal effort and higher maternal GC levels will be associated with lower rates of social play, less behavioral independence, and slower growth. Mothers who experienced early life adversity themselves or mothers that currently experience adverse conditions will have infants with lower rates of social play, less behavioral independence, and slower growth than other females.
1b. Alternatively, signals of low maternal capital might lead infants to prioritize growth over behavioral development: If so, higher levels of maternal effort and higher maternal GCs will lead to lower rates of social play, less behavioral independence, and faster growth.
2. Infants face tradeoffs: There will be a negative relationship between rates of play and somatic growth, particularly for infants born to mothers with fewer resources because these infants are operating under greater constraints. For behavioral development, we focus on play because of its high energetic costs.
3. Previous research indicates that developmental trajectories and tradeoffs may differ between male and female infants. For example, during periods of high food availability, female Assamese macaques prioritize growth and males prioritize play (Berghänel et al., 2015), likely a reflection of sex-based differences in the developmental traits most influential for reproductive success and fitness. In species in which male development is more energetically costly than female development, and in species with greater male reproductive skew, males are expected to be more sensitive to signals of resource availability to minimize mortality risk during poor conditions and maximize response when resources are great (Clutton-Brock et al., 1985). Thus, we will explore the possibility that males respond differently than females to variation in maternal signals and characteristics.

## Methods

### Study site and subjects

We studied four groups of wild baboons that range on the eastern Laikipia Plateau of central Kenya. These groups are monitored by the Uaso Ngiro Baboon Project (UNBP), directed by Dr. Shirley Strum. The study groups range in an area that is topographically diverse and is located 1718m above sea level. The habitat is dry savanna with grassy plains, acacia woodlands, and riverine woodlands on the edge of dry sandy rivers. Annual rainfall is typically concentrated in two wet seasons (March-June, November-December; (Barton, 1993), though droughts are increasingly common). *Opuntia stricta*, an invasive non-indigenous cactus, has become an important part of the diet for all of the groups monitored by the UNBP (Strum et al., 2015). Access to the *Opuntia stricta* fruit has reduced seasonal variability in food availability and shortened interbirth intervals (Strum, unpublished data). Three of the study groups PHG, ENK, and YNT occupied overlapping home ranges and the fourth study group, NMU, ranged in a different area. Individuals in PHG, ENK, and YNT had more *Opuntia stricta* in their diet than those in NMU. Demographic records span the entire study period from the oldest mother’s birth in 1998 till 2017. Observers update demographic records daily and record births, deaths, and disappearances. Maternal kinship relationships among natal females were known from genealogical records extending back to the early 1970s.

Observers collected data on the local ecology and acute environmental challenges. Data on herbaceous biomass are collected each month using the slanting pin intercept technique angled 65 degrees from vertical (McNaughton, 1979) and converted into biomass in gr/m2 using the adjusted equation HB=total hits X 0.847 (McNaughton, 1979; Western & Lindsay, 1984). Encounters with humans and unfamiliar baboons represent acute challenges. During 2016-2017, we recorded 3 infant deaths and 1 adult death due to human-baboon interactions. The presence of unfamiliar individuals from other groups is associated with higher GCs and increased risk of wounding in several primate species (baboons: (Beehner et al., 2005; MacCormick et al., 2012); gelada monkeys (*Theropithecus gelada*): (Schneider-Crease et al., 2020). We recorded all encounters with humans and unfamiliar baboons *ad libitum*. We counted as acute challenges the total number of encounters with humans and unknown male baboons each month.

### Maternal characteristics

#### Behavioral observations (observation period: 2016-2017)

Observers conducted approximately 2106 complete 15-min focal samples from October 2016 to December 2017 on 40 focal infants (ages: birth to 1 year). Each of the 40 focal offspring was observed on average 9.2 times per month (range: 1-19 times/month). During focal samples, observers recorded activity state, social interactions, and vocalizations on a continuous basis (Altmann, 1974). All behavioral data were collected on hand-held computers (Palm Zire 21) in the field and later transferred onto computers for error checking and storage in the NS Basic program. In addition to focal data on infants, data on agonistic contests were collected from adults and subadults by UNBP observers to establish monthly dominance ranks. We calculated maternal rank relative to the total number of females in the hierarchy (Levy et al., 2020), such that ranks range from 0 to 1 and higher numbers indicate higher rank.

During focal follows, nursing and carrying were recorded as durational states. Maternal behavioral effort was calculated as the proportion of observation time spent nursing offspring and the proportion of time spent carrying offspring (Altmann & Samuels, 1992; Ross, 2001). For each day of focal observation, we calculated the total amount of time that offspring spent nursing (i.e. time with nipple in mouth) and being carried by their mother and divided this by the total amount of time observed.

#### Fecal Collection, Hormonal Extraction, and Hormone Assays (observation period: 2016-2017)

We included a total of 552 fecal samples from the 38 lactating mothers in this study (average=3.03 samples per mother per month). The protocol for collection, extraction, and storage have been validated and described in detail in primates (Beehner & McCann, 2008). Within 10 minutes following deposition, the fecal sample was mixed thoroughly with a wooden spatula, and an aliquot of the mixed sample (~ 0.5 g wet feces) was placed in 3 mL of a methanol/acetone solution (4:1). The solution was immediately homogenized using a battery-powered vortex. The weight of the dry fecal matter was later determined using a battery-powered, portable scale to ± 0.001 g. Approximately 4–8 h after sample collection, 2.5 mL of the fecal homogenate was filtered through a 0.2 μm polytetrafluoroethylene (PTFE) syringeless filter (Fisher cat #09-921-13), and the filter was then washed with an additional 0.7 mL of methanol/acetone (4:1). We then added 7 mL of distilled water to the filtered homogenate, capped and mixed the solution, and loaded it onto a Waters Corp Sep-Pak reverse-phase C18 solid-phase extraction cartridge (Fisher cat #50-818-645). Prior to loading, Sep-Pak cartridges were prepared according to the manufacturer’s instructions (with 2 mL methanol followed by 5 mL distilled water). After the sample was loaded, the cartridge was washed with 2 mL of a sodium azide solution (0.1%). All samples were stored on cartridges in separately sealed bags containing silica beads. Cartridges were stored at ambient temperatures for up to 10 days, after which all samples were stored at −20 °C until transported at room temperature to Arizona State University for analysis. In the laboratory, steroids were eluted from cartridges with 2.5 mL 100% methanol and subsequently stored at − 20 °C until the time of enzyme immunoassay (EIA).

We analyzed GCs in our samples using a group-specific EIA for the measurement of immunoreactive 11β-hydroxyetiocholanolone (Frigerio et al., 2004), which has been used to monitor glucocorticoids in other primate species and validated biologically with an ACTH challenge test in olive baboons (Higham et al., 2009). Samples (1:60 fecal extract to buffer) and standards were added to each plate in duplicate (50 uL/well), followed by 50 uL of biotin-labeled hormone and 50 uL of antibody to each well. Plates were incubated for at least 18 h at 4°C, and no more 24 hours. Plates were washed with a wash solution (PBS solution with 0.05% tween) and 150 uL of streptavidin-peroxidase was added to each well, incubated for one hour, and then the plate was washed again. We added 100 uL of TMB substrate solution to each well. Plates were incubated while shaking for 55-60 mins and the reaction was stopped with the addition of 50 uL of sulfuric acid. The plate was read at wavelength of 450 nn on a Synergy H2 plate reader. Cross-reactivities for the 11β-hydroxyetiocholanolone assay include 5β-androstane-3α-ol-17-one, 11-oxo-etiocholanolone and <0.1% for corticosterone, cortisol, 5α-androstane-3,11,17-trione, 5β-androstane-3,11,17-trione, testosterone, 5α-androstane-3,17-dione, 5β-androstane-3,17-dione, androstendione, 5β-androstane-3β-ol-17-one, 5β-androstane-17-one, dehydroepiandrosterone, and androsterone; these are characterized in: Ganswindt et al, 2003. Serially diluted fecal extracts were parallel to the standard curve (F = 0.10, p=0.77). We used low (avg: 15.9 ng/g) and high (avg: 53.8 ng/g) concentrations of pooled baboon samples as controls. Inter-assay CVs were 18.6% and 24.4% respectively and intra-assay CVs were 7.7% and 11.4%. We did not have maternal fecal GC samples to pair with each infant focal follow, so GC samples were averaged for each month of observation.

#### Assessment of Mothers’ Early Life Adversity (long-term data: 1998-2015)

Following Patterson et al 2021, we considered 5 measures of early life adversity for the mothers that we studied:

(1) Herbaceous biomass was used as an indicator of drought-like environmental conditions. We recorded monthly biomass data separately for two ranging areas. NMU troop occupied one ranging area and PHG, ENK, and YNT occupied the other range. Biomass was averaged for the year of each mother’s birth with lower biomass inverted to indicate a higher adversity score. (2) Group size at birth was used as an indicator of the extent of within-group competition. Group size was defined as the number of adult and subadult males and females in the troop on the day the mother was born. (3) Maternal loss was defined as the age at which the mother lost her own mother. This score was then inverted so that maternal loss at an earlier age was associated with a higher adversity score. We include maternal loss after the period of nutritional independence because maternal death after weaning continues to have substantial effects on offspring survival and fitness(Crockford et al., 2020; Foster et al., 2012; Nakamura et al., 2014; Samuni et al., 2020; Stanton et al., 2020; Zipple et al., 2021). We use 4 years of age as a cutoff because we are interested in early life experiences and 4 years marks the earliest age at menarche in several cercopithecine populations (Patterson et al., 2024; Strum & Western, 1982; Tung et al., 2016). Mothers who lost their own mother after the age of 4 years received a zero for this component of early life adversity. (4) Maternal interbirth interval (IBI) was measured as the time between the mother’s own birth to the birth of her next younger sibling, an indicator of her own mother’s condition and ability to invest. We consider longer investment periods to represent greater adversity for several reasons. First, longer IBIs are associated with older age and lower rank suggesting a link to poor maternal condition (reviewed in (Harcourt, 1987)). Second, in this population larger group size at birth is associated with longer IBIs (Patterson et al., 2021). Third, the spread of *Opuntia stricta* lowered IBIs substantially (UNBP unpublished data). Shorter IBIs may also be associated with costs, but here, we follow previous methodologies used in this study population which found that longer IBIs had similar effects as other forms of adversity on maternal effort and glucocorticoid levels (Patterson et al., 2021). (5) Primiparity was measured as whether the mother was a first-born (Pittet & Hinde, 2023). We used continuous measures for all components of early life adversity except primiparity. All of the continuous measures were normalized so values range from zero to one and can be summed to create a cumulative score. Primiparity was scored as 1 for first borns, and 0 for non-first borns. All five scores were summed to create the cumulative adversity index.

We also measured how long mothers had access to *Opuntia stricta*. Animals in PHG, ENK, and YNT started to eat *O. stricta* fruit regularly in 2000 and animals in NMU started to eat it regularly in 2012. We calculated how old each mother was when her troop started to consume *O. stricta.* Age zero was used if *O. stricta* was already present at her birth.

### Infant outcomes (observation period: 2016-2017)

#### Infant behavior

During focal follows, all approaches to within 1m and departures out of 1m between mother and offspring were recorded. Offspring independence was calculated as the number of departures (1m distance) initiated by the offspring away from their mother each day, controlling for the time mother and offspring were in proximity that day. Social play involving physical contact with conspecifics (play, hereafter) was recorded continuously (Fairbanks, 2000). For each day of focal observation, we counted the number of play bouts in which a focal participated.

#### Infant size and growth trajectories

We used digital photogrammetric methods to measure infant body size methods. We mounted parallel lasers that were 4cm apart to a Nikon DSL camera. (blueprints: https://amylulab.weebly.com/labwork.html.) The laser projections create a measurement scale within each photograph to assess body size. We used the scale provided by the parallel lasers to measure shoulder-rump (SR) length. Photographs were taken while infants were standing quadrupedally on level ground with the longitudinal axis of their body perpendicular to the photographer. To measure SR length, we follow previous studies (Lu et al., 2016; Rothman et al., 2008) and use the following formula:

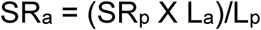

Where SR represents the distance between the shoulder and rump, L represents the distance between the laser points, the subscript *a* represents actual distance, and the subscript *p* represents the distance in the photograph.

All photographs were measured by SKP and at least one of three other measurers (KP, BL, SK). We made 884 measurements of 281 photos. We took the mean of multiple measures for each photograph. The mean between-measurer CV within photos was 2.7%. There were 71 cases in which we obtained more than one photograph of an infant on the same day. The mean between photograph CV was 3.3%. There were 195 daily mean body size estimates. We removed one outlier. To mitigate the consequences of potential error in body size estimates, we only included infants for whom we had at least two body size estimates that were taken at least 30 days apart. The final growth dataset includes 21 infants (2-23 photos per infant; mean=8.8 photos per infant).

### Statistical Modeling

We fit Bayesian models using Hamilton Markov chain Monte Carlo (MCMC) with r-STAN v.2.18.2 (Stan Development Team 2018) in R v.3.3.2 (R Core Team, 2021) using the map2stan function in the ‘rethinking’ package v.1.59 (McElreath, 2016). All continuous predictors were transformed to a mean of 0 and a standard deviation of 1. We used conservative, regularizing priors on all of our predictor parameters to ensure that our models were skeptical of large effects. It is difficult to use model parameters to perceive joint effects on posterior predictions so we rely on graphs of model predictions over raw data to assess the patterns of results. These figures provide information regarding the relative magnitude and certainty of the effects of variables of interest on the scale of the outcome variable. Model code for this dataset can be found here: https://github.com/skpatter/maternaleffects.

#### Models of offspring behavior

To determine what factors predicted infant play and departures from mother, we fit zero-inflated Poisson (ZIP) models. The outcome variable (count of play and count of leaves) is predicted by mixing Bernoulli and Poisson components of the model. Positive coefficients from the Bernoulli component mean that the offspring is less likely to play or leave, and positive values for the Poisson component indicate higher counts of behavior conditional upon the behavior occurring at all. Their joint likelihood is calculated by multiplying the likelihoods of the Bernoulli and Poisson outcomes.

We modeled offspring behavior as a function of maternal signals (proportion of observation time spent nursing, proportion of observation time spent carried by mother, maternal monthly mean GCs). We included several covariates to control for infant age, infant sex, current social and ecological conditions (current monthly group size, current monthly acute challenges, and current monthly herbaceous biomass) and maternal capital (early life adversity score of mother, parity of mother, rank of mother, presence of opuntia at mother’s birth) We also included an offset for observation time to estimate the rate of play and rate of departures. When modelling infant departures from mother, we controlled for the total time the infant and mother were within 1m proximity. We ran the play and independence models with and without a quadratic term for offspring age. The models with the quadratic term received full model weight for both outcomes so this term is included in the final models. We included varying effects for infant ID. We ran two versions of the play and independence models: 1) the first model did not include interaction terms, and 2) the second model included an interaction between infant sex and nursing, carrying, and maternal GCs (i.e., sex*nursing, sex*carrying, sex*GCs). Model comparisons are reported in the Results.

#### Models of offspring growth

We ran four initial models to generate the variables needed for our main growth model. These variables are growth rate, and mean nursing level, mean carrying level, and mean GCs that account for infant age. The first initial model was built to estimate individual growth rates. We modeled body size as a function of infant age, fit with a Gaussian model. Infant ID was treated as a varying intercept and a varying slope multiplied by age, allowing for an individual slope (i.e., growth rate) for each infant. We used linear estimates for growth because we only examined individuals during infancy when growth appears linear (Figure S1). The model produced mean intercepts and slopes with offsets for each infant. As expected, infant age had a positive relationship with body size (B = 0.98, SD=0.05). The next three models were built to calculate a nursing, carrying, and maternal GC estimates to be paired with each infant’s growth rate. The models with the proportion of observation time spent nursing and carrying were fit with gamma distributions because the values were skewed right and constrained to be positive. We included infant age as a control variable and a varying intercept for ID. For maternal GCs, we used a Gaussian model with infant age and a varying intercept for ID. Age had a negative relationship with both nursing time (B = −0.27, SD=0.02) and carrying time (B = −0.27, SD=0.02). GCs increased across lactation (B = 0.16, SD=0.04). We extracted the individual varying intercepts for nursing, carrying, and GCs to use in the model estimating growth rates.

We fit Gaussian models to determine what factors predicted growth rates. Growth rates were estimated by the individual varying slopes from the first initial model described above. As predictor variables, we include the varying intercept outputs from the initial models for nursing, carrying, and maternal GCs. As covariates, we also included infant sex, and factors associated with maternal capital (maternal early life adversity, age at mother’s introduction to *Opuntia*, parity) and current social and ecological conditions (mean maternal rank for the observation period, and mean herbaceous biomass for the observation period). There was a high correlation among the current environmental variables for this model due to the use of mean values across the observation period, and we thus chose to include current herbaceous biomass rather than current group size or current challenges because food availability is a biologically meaningful predictor of growth rates. We ran two versions of the final growth model: 1) the first model did not include interaction terms, and 2) the second model included an interaction between infant sex and nursing, carrying, and maternal GCs (i.e., sex*nursing, sex*carrying, sex*GCs). Model comparisons are reported in the Results.

#### Tradeoff Model

To test for a tradeoff between play and growth, we reran the play model with one additional fixed effect: growth rate. To explore our prediction that a tradeoff will be more pronounced among offspring born to mothers with behavioral and physiological indicators of potentially lower maternal capital (high GCs, more nursing and carrying), we ran two models: 1) the first model did not include interaction terms, and 2) the second model included an interaction between infant growth rate and nursing, carrying, and maternal GCs (i.e., growth*nursing, growth *carrying, growth *GCs). Model comparisons are reported in the Results.

## Results

### Offspring play behavior

Maternal effort and GC levels influenced the frequency of infant play. The model without an interaction between offspring sex and nursing time, carrying time, and maternal GCs received full weight. Offspring who spent more time nursing and being carried played less than offspring who spent less time nursing and being carried by mother (Fig. 1, Table S1). Offspring born to mothers with higher GCs during lactation also played less than offspring of mothers with lower GCs (Fig. 1, Table S1).

**Figure 1.**
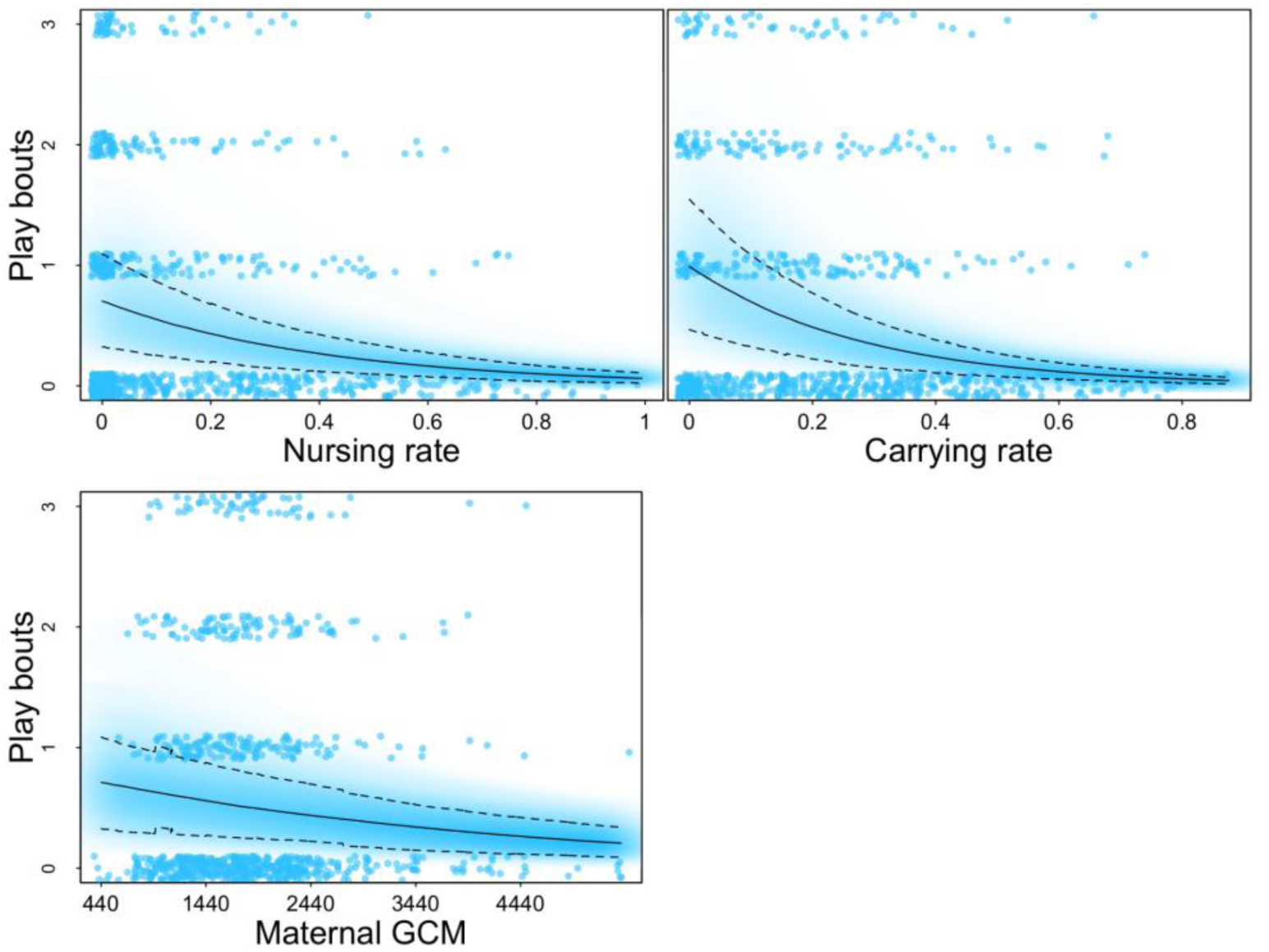
Offspring play behavior Model averaged posterior predictions for the influence of nursing, carrying, and maternal GCs on the number of bouts of play. The solid line represents the mean estimate. The dashed lines represent the 89% highest posterior density interval. The blue cloud shows the full posterior predictions, with darker areas representing higher densities. Model sample sizes are as follows: 40 infants, 38 mothers, and 1002 data points.

Both maternal early life conditions and current conditions influenced the frequency of offspring play. Higher levels of maternal early life adversity were associated with less offspring play (Fig. S2). Current herbaceous biomass had a strong positive effect on the rate of play bouts (Fig. S6), while current acute challenges had a negative effect on play (Fig. S7). Mothers who lived longer prior to the introduction of *Opuntia* produced offspring who played more compared to mothers who had access to the fruit for all or most of their lives (Fig. S3). Maternal rank, maternal parity, and current group size did not have substantial effects on the rate of offspring play (Figs. S4, S8, S9). Offspring age had a nonlinear effect with play increasing until around 4 months of age and then slowly declining (Fig. S5). Sons played at higher rates than daughters (Fig. S10).

### Offspring independence

The model with an interaction between offspring sex and nursing time, carrying time, and maternal GCs received full weight. Offspring who spent more time nursing were less independent (i.e., departed from their mother less often) than offspring who nursed less (Fig. 2, Table S2). Offspring born to mothers with higher GCs during the lactation period showed less independent behavior than offspring born to mothers with lower GCs, but this pattern was only exhibited by sons (Fig. 2, Table S2). Carrying time was not a strong predictor of independent behavior for either sex (Fig. 2, Table S2).

**Figure 2.**
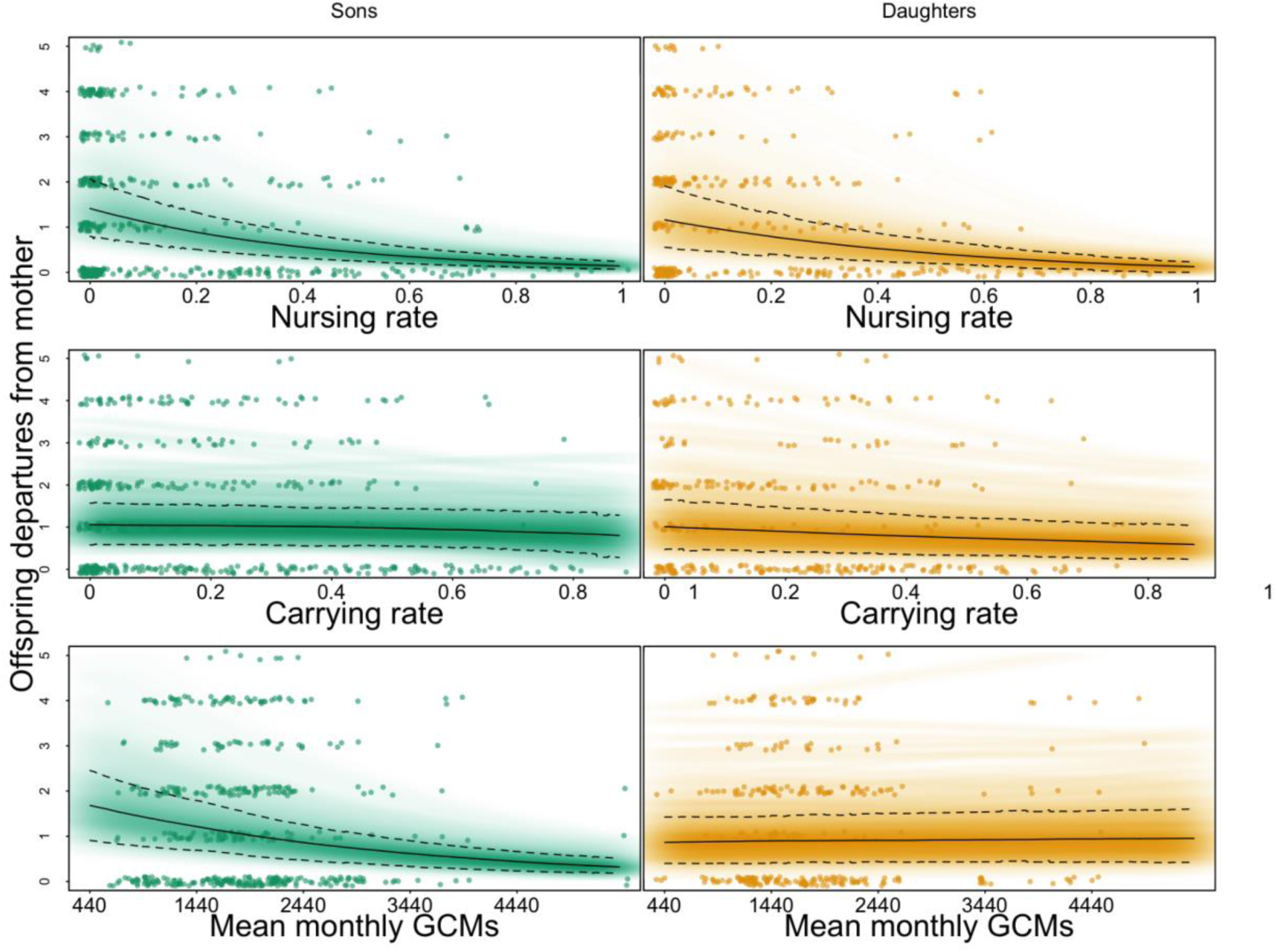
Offspring independence behavior Model averaged posterior predictions for the influence of nursing, carrying, and maternal GCs by offspring sex on the number of offspring departures from their mothers. The solid line represents the mean estimate. The dashed lines represent the 89% highest posterior density interval. The green (sons) and yellow (daughters) clouds show the full posterior predictions, with darker areas representing higher densities. Model sample sizes are as follows: 40 infants (23 sons, 17 daughters), 38 mothers, and 1002 data points.

More herbaceous biomass predicted more independent behavior (Fig. S14), whereas more current acute challenges predicted less independent behavior (Fig. S15). The other predictors of offspring independence were relatively uncertain. Infants whose mothers experienced more early life adversity were slightly less independent than those born to mothers who had less early life adversity (Fig. S11). Offspring born to higher ranking mothers were also slightly less independent than those born to lower ranking mothers (Fig. S12). Mothers who lived longer prior to the introduction of *Opuntia stricta* produced offspring who were less independent compared to mothers who had access to the fruit for all or most of their lives (Fig. S16). Sons were slightly more independent than daughters, but the distributions mostly overlap (Fig. S18). There was no clear effect of parity (Fig. S17).There was a nonlinear effect of offspring age with departures from mother increasing from birth till 2-3 months old and then decreasing with age (Fig. S13).

### Growth

The model without an interaction between offspring sex and nursing time, carrying time, and maternal GCs received almost full weight. Offspring whose mothers spent more time nursing and carrying them grew more slowly than offspring whose mothers nursed and carried them less, though the model was relatively uncertain about the nursing effects (Fig. 3, Table S3). Maternal GCs during the lactation period had a slight positive association with offspring growth rates but the model was relatively uncertain about this association (Fig. 3, Table S3).

**Figure 3.**
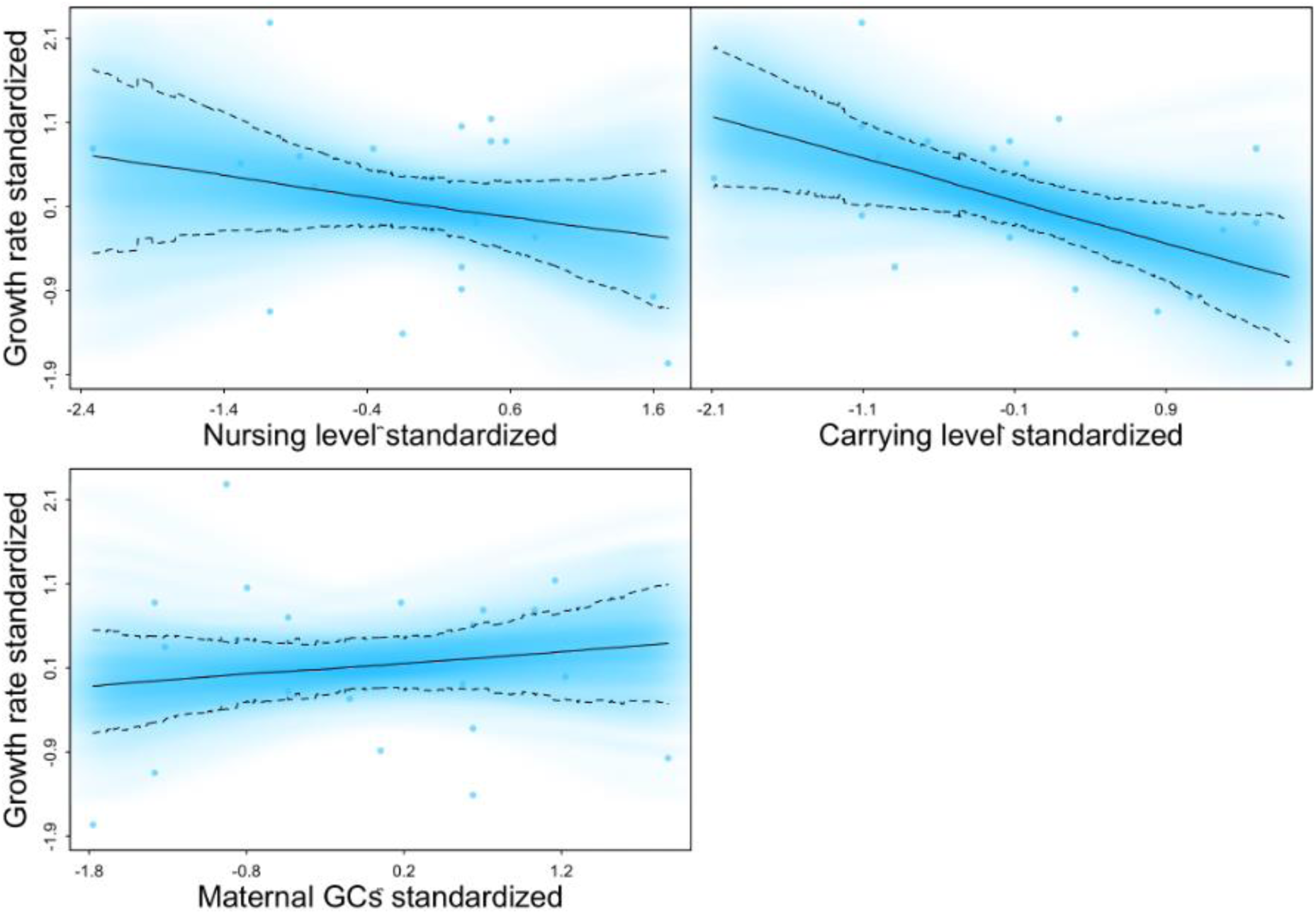
Offspring growth rates Model averaged posterior predictions for the influence of nursing, carrying, and maternal GCs on offspring growth. The solid line represents the mean estimate. The dashed lines represent the 89% highest posterior density interval. The blue cloud shows the full posterior predictions, with darker areas representing higher densities. Model sample sizes are as follows: 21 infants, 21 mothers, and 21 data points.

The growth model was also suggestive but uncertain about the following effects. Offspring born to mothers who experienced more early life adversity grew faster than offspring born to mothers with less early life adversity (Fig. S19). Mothers who lived longer prior to the introduction of *Opuntia* produced offspring who grew slightly faster than mothers who had access to the fruit for longer (Fig. S20). Offspring born to higher ranking and multiparous mothers grew faster than offspring born to lower ranking and primiparous mothers (Fig. S21, Fig. S24). Greater herbaceous biomass was associated with faster growth rates (Fig. S22). Daughters grew faster than sons on average, but the distributions were mostly overlapping (Fig. S23).

### Tradeoff between play and growth

Because play is energetically costly, we expected to find a tradeoff between play and growth. Overall, offspring who grew faster played less than offspring who grew at slower rates (Table S4). The model with interactions between offspring growth and nursing rates, carrying, and maternal GCs received full weight. Among offspring who nursed at lower rates and offspring born to mothers with lower GCs, there is a negative association between growth and play (Fig. 4, Table S4). Among offspring who nursed at higher rates and offspring born to mothers with higher GCs, this relationship between growth and play was diminished (Fig. 4, Table S4). It appears that offspring who nursed at higher rates and were exposed to higher maternal GCs exhibited low levels of play regardless of growth rate, whereas offspring who nursed at lower rates and were exposed to lower maternal GCs played at higher rates if they grew more slowly. There was no interaction between carrying and growth.

**Figure 4.**
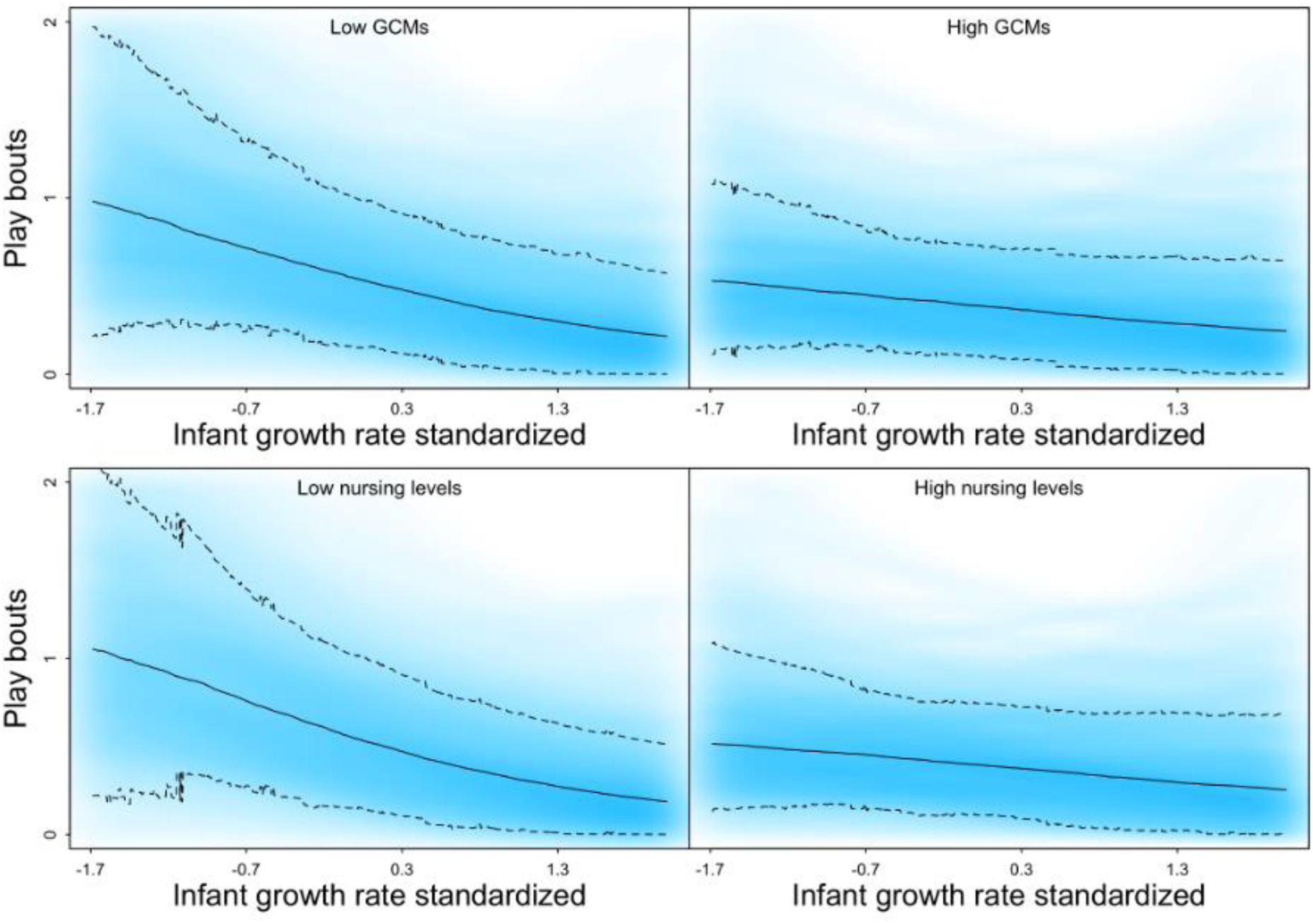
Offspring play versus growth Model averaged posterior predictions for the influence of offspring growth on play rates and the interactions with nursing levels and maternal GCs. The solid line represents the mean estimate. The dashed lines represent the 89% highest posterior density interval. The blue cloud shows the full posterior predictions, with darker areas representing higher densities. Model sample sizes are as follows: 22 infants, 22 mothers, and 676 data points.

## Discussion

The findings of this study suggest that maternal behavioral care and physiological signals guide how offspring navigate development. Higher maternal effort and maternal GCs across lactation were associated with reduced rates of play and fewer departures from the mother. A mother’s own early life adversity had an independent negative effect on offspring play and independence. Low ranking mothers and mothers who had access to the novel *Opuntia stricta* fruit for a shorter period of their lives produced offspring that were less independent. These positive links between maternal condition and behavioral development are in line with similar patterns in other primate species (blue monkeys: (Förster & Cords, 2005); rhesus macaques: (Hinde et al., 2015; Mccormack et al., 2006); yellow baboons: (Nguyen et al., 2012); but see vervet monkeys: (Fairbanks & Hinde, 2013)), and contrasts with those in rodents (reviewed in (Catalani et al., 2011)). Together these patterns suggest infants born into a maternal environment with fewer resources allocate fewer resources to behavioral development. Such a developmental trajectory might carry the cost of slower motor skill development (Belding’s ground squirrels: (Nunes et al., 2004); Assamese macaques (*Macaca assamensis*): (Berghänel et al., 2015)) and poor fitness outcomes (age at dispersal & reproductive success in squirrels: (Nunes et al., 2004); survival in horses: (Cameron et al., 2008)).

Findings on growth patterns in this study were complex. Greater maternal effort was associated with slower offspring growth as predicted (prediction 1a). This relationship is consistent with the idea that adverse conditions constrain energy availability and reduce growth (Lu et al., 2019). However, higher maternal GCs during lactation and maternal early life adversity predicted faster growth as expected under our alternative prediction (1b). Given that maternal early life adversity, low maternal rank, and current challenges predict more maternal effort, and maternal early life adversity and lower herbaceous biomass predict higher maternal glucocorticoids in this population (Patterson et al., 2021), there is likely a complex interplay among maternal early and current life experiences, maternal behavioral and physiological signals, and impacts on offspring development. All of these variables were included in the statistical models to examine how the different components shape offspring growth, but future work would benefit from disentangling the causal pathways. While our models were uncertain about the effects of GCs and maternal adversity on growth, these patterns are similar to previous studies, suggesting higher GCs could instruct infants to allocate more energy into growth at the expense of other dimensions of development (Allen-Blevins et al., 2015; Hinde et al., 2015). As noted earlier, the timing of maternal GC signals and dynamic changes in the signals over the period of care lead to different growth trajectories in rhesus macaques (Hinde et al., 2015; Petrullo et al., 2019). In humans, milk GCs at 3 months and maternal salivary GCs were negatively associated with infant growth (Hahn-Holbrook et al., 2016; Thayer et al., 2012), but several other studies found no association between milk GCs and infant growth (Feizabad et al., 2025; Hollanders et al., 2019; Zielinska-Pukos et al., 2022). How these complex processes contribute to our results is particularly hard to disentangle because we collected fecal GCs rather than cortisol in mother’s milk.

Our findings suggest maternal behavioral care and physiological signals guide infant priorities in the tradeoff between somatic growth and play behavior during maternal dependence. There was a negative relationship between the rate of play and growth rate as was previously demonstrated in wild Assamese macaques (Berghänel et al., 2015, 2016). Interestingly, the tradeoff was more pronounced among offspring exposed to lower maternal GCs and lower nursing levels. We originally predicted that infants developing under greater energetic constraints would display more intense tradeoffs, whereas less constrained infants might be able to invest in both behavioral development and fast growth. Instead, our results suggest that offspring exposed to lower nursing levels and lower maternal GCs are moderately constrained and can prioritize either somatic growth or behavioral development. Offspring exposed to higher nursing levels and higher maternal GCs are more severely constrained and seem to prioritize growth over behavioral development if they have available resources to invest. Based on these results, we distinguish among three levels of constraints: (1) no constraints and no trade-offs because offspring can invest in both growth and behavior, (2) mild constraints and trade-offs such that offspring can invest in one or the other more, and (3) major constraints and no trade-offs because offspring can only invest in growth. Building off these findings, it may be fruitful for future work to investigate the potential tradeoffs among growth, behavior, and immune health. Milk GCs and other milk bioactives may play a role in shaping tradeoffs among these three dimensions. In humans, milk GCs are associated with milk antibodies and proinflammatory responses, which in turn shape risk of infant illness and immune development (Anyim et al., 2023; Matyas et al., 2024; Rosen-Carole et al., 2024).

External environmental factors also affected infant development. Although mothers are predicted to buffer offspring to some extent from the external environment, this protection will lessen and effects of external environments are expected to increase as infants develop and increase their locomotive and nutritional independence (reviewed in (Wells, 2010, 2014, 2019)). Current herbaceous biomass had a strong positive effect on the rate of play bouts and independent behavior, and an uncertain but positive effect on growth, which is consistent with previous work linking low food availability to lower rates of play, slower growth, and smaller body size in a range of species (Altmann & Alberts, 2005; Berghänel et al., 2015; Bock & Johnson, 2004; Cameron et al., 2008; Espinosa et al., 1992; Hinde, 2013; Johnson, 2003; Lee, 1983; Li & Rogers, 2004; Lochmiller et al., 2000; Martin & Caro, 1985; McAdam & Millar, 1999; Muller-Schwarze et al., 1982; Nunes et al., 2004; Sharpe et al., 2002; Siviy & Panksepp, 1985; Stone, 2008; Strum, 1991; Sugiyama & Ohsawa, 1982). As expected, in response to frequent encounters with humans and strange adult male baboons, infants departed from their mothers less and played at lower rates. Effects of the external environment are likely due to a combination of direct effects on the infant and indirect effects through its influence on maternal physiology and behavior (Beehner et al., 2005; Beehner & McCann, 2008; Gesquiere et al., 2011; Schneider-Crease et al., 2020).

Sons and daughters navigated development in somewhat different ways. Sons played at higher rates, were more independent, and on average, grew more slowly than daughters. Our findings parallel previous studies in a range of primate species that found male infants play more and exhibit more behavioral independence from their mothers than female infants (reviewed in (Lonsdorf, 2017; Meredith, 2013)). Sons and daughters also responded differently to maternal physiology with regard to their independence behavior, but not play or growth. In response to elevated maternal GCs, sons exhibited substantially lower independence while daughters did not adjust behavior, thus contributing to the limited research in primates that indicates sons may be more sensitive to maternal GC signals than are daughters, and likely have different timepoints and durations of sensitivity (Hinde et al., 2015; Murray et al., 2018; Petrullo et al., 2019). However, there are also cases in which female infants are more sensitive (Dettmer et al. 2018; Hinde et al 2015). For example, infant female rhesus macaques are sensitive to absolute levels of milk GCs, and infant males are more sensitive to dynamic changes in milk GCs across lactation (Hinde et al., 2015). Our findings lend support to the notion that there are differences in the ways sons and daughters prioritize resources and energy (Brown et al., 2024; Hinde et al., 2015; Petrullo et al., 2019). Females seem to prioritize growth while males prioritize play and motor skill development. Similar patterns have been documented in Assamese macaques (Berghänel et al., 2015).

Our study had several limitations. First, we used maternal fecal GCs as a proxy for milk GCs, which are directly transmitted to the offspring. Although this measure is indirect, the use of maternal fecal GCs provides an overview of maternal biology across lactation rather than sampling at constrained and fewer timepoints, which must be done with milk sampling. Second, we used photogrammetry to monitor body size and growth during the first year of life, however, it is very difficult to obtain a large number of usable photographs of infants posing in the proper position and orientation; thus, our sample of usable photographs was small and there was considerable uncertainty in estimates of the effects of variables linked to infant growth. Although this method involves measurement error, photogrammetry provides a noninvasive option when trapping animals is not feasible. Finally, while we demonstrate associations between the maternal environment and offspring development, some portion of these effects may be explained by shared genes between mother and offspring. Unfortunately, we are unable to measure and account for the effect of genetics in our analyses. We encourage future studies with larger sample sizes to use statistical tools such as the “animal model” to incorporate pedigrees or genetic relatedness to estimate both maternal genetic and nongenetic effects (Brent et al., 2017; Kruuk, 2004; Patterson et al., 2024; Wilson et al., 2010).

In conclusion, maternal behavioral and physiological signals influenced how infants navigated multiple dimensions of development. Mothers with lower capital, or a lower capacity to invest, produced offspring who were more constrained as they navigated development, generally exhibiting reduced somatic and behavioral development but prioritizing allocation to growth over behavior when resources allowed. Future studies exploring the longer-term consequences of the ways infants use maternal signals to orchestrate developmental trajectories are needed to better understand the evolution of these processes and their fitness consequences.

## Acknowledgements

We thank the Office of the President of the Republic of Kenya and the Kenya Wildlife Service for permission to conduct this field research. This work was supported by the National Science Foundation (BCS-1732172; GRFP-1841051), the Leakey Foundation, and Arizona State University. We thank Kate Abderholden, Megan Best, Megan Cole, Moira Donovan, Alexandra Duchesneau, Jessica Gunson, Bailey Lindquist, Molly McEntee, Kapulani Patterson, Laura Peña, Eila Roberts, Nia Stratton, Sinera Williams, and Leah Worthington for their contributions to data collection. We thank the staff of the Uaso Ngiro Baboon Project, particularly Jeremiah Lendira, James King’au, Joshua Lendira, and Frances Molo for their help and companionship in the field; David Muiruri for invaluable assistance with logistics and data management; and the African Conservation Centre for facilitating the UNBP project and assisting us with our work.

## Notes

### Competing Interest Statement

The authors have declared no competing interest.

