## Supplementary Materials for "Maternal glucocorticoids and behavior shape offspring developmental trade-offs in wild baboons"

Table S1.Coefficients for model evaluating the effect of maternal effort and maternal GCs on offspring social contact play

| | p component | | l component | |
|---|---|---|---|---|
| | Mean | StdDev | Mean | StdDev |
| Intercept | -2.85 | 0.60 | 0.46 | 0.24 |
| Nursing | -0.33 | 0.27 | -0.48 | 0.05 |
| Carrying | 0.02 | 0.27 | -0.67 | 0.05 |
| GCs | -0.31 | 0.20 | -0.20 | 0.03 |
| ELA | -0.38 | 0.39 | -0.18 | 0.14 |
| Opuntia | -0.48 | 0.35 | 0.01 | 0.12 |
| Rank | -0.28 | 0.30 | -0.07 | 0.11 |
| Parity | -1.19 | 0.67 | -0.17 | 0.27 |
| Sex | 0.83 | 0.59 | -0.35 | 0.22 |
| Infant age | -0.66 | 0.24 | -0.40 | 0.05 |
| Infant age sq | 0.54 | 0.19 | -0.14 | 0.04 |
| Group size | 0.07 | 0.42 | -0.07 | 0.15 |
| Challenges | -0.64 | 0.26 | -0.22 | 0.04 |
| Biomass | 0.53 | 0.22 | 0.32 | 0.04 |

Table S2. Coefficients for model evaluating the effect of maternal effort and maternal GCs on offspring independent behavior (departures from mother)

| | p component | | l component | |
|---|---|---|---|---|
| | Mean | StdDev | Mean | StdDev |
| Intercept | -4.37 | 0.80 | 0.89 | 0.34 |
| Nursing | -0.46 | 0.37 | -0.46 | 0.05 |
| Carrying | 0.85 | 0.31 | -0.01 | 0.04 |
| GCs | -0.25 | 0.35 | -0.27 | 0.03 |
| Nursing*Sex | 1.16 | 0.56 | 0.09 | 0.07 |

| | | | | |
|---|---|---|---|---|
| Carrying*Sex | -0.64 | 0.48 | -0.09 | 0.05 |
| GCs*Sex | -0.23 | 0.46 | 0.28 | 0.04 |
| ELA | -0.56 | 0.50 | -0.16 | 0.18 |
| Opuntia | 0.31 | 0.41 | -0.20 | 0.15 |
| Rank | -0.51 | 0.38 | -0.16 | 0.14 |
| Parity | -0.46 | 0.89 | -0.07 | 0.38 |
| Sex | 0.09 | 0.77 | -0.15 | 0.29 |
| Infant age | -0.64 | 0.32 | -0.56 | 0.04 |
| Infant age sq | 1.32 | 0.21 | -0.10 | 0.03 |
| Group size | 0.37 | 0.55 | 0.32 | 0.21 |
| Challenges | -0.48 | 0.26 | -0.27 | 0.03 |
| Biomass | -0.05 | 0.30 | 0.14 | 0.03 |

Table S3. Coefficients for model evaluating the effect of maternal effort and maternal GCs on offspring growth rate

| | Mean | StdDev |
|---|---|---|
| Intercept | -0.25 | 0.42 |
| Nursing Levels | -0.28 | 0.31 |
| Carrying Levels | -0.52 | 0.25 |
| Maternal GCs | 0.14 | 0.21 |
| Maternal ELA | 0.17 | 0.35 |
| Maternal Opuntia | 0.06 | 0.30 |
| Maternal Parity | 0.38 | 0.52 |
| Maternal Rank | 0.25 | 0.27 |
| Infant Sex | 0.23 | 0.54 |
| Biomass | 0.19 | 0.31 |

Table S4. Coefficients for model evaluating the relationship between offspring growth and play bouts

| | p component | | l component | |
|---|---|---|---|---|
| | Mean | StdDev | Mean | StdDev |
| Intercept | -3.89 | 0.85 | 0.09 | 0.38 |
| Nursing | 0.20 | 0.32 | -0.30 | 0.06 |
| Carrying | -0.09 | 0.30 | -0.82 | 0.06 |
| GCs | -0.35 | 0.22 | -0.31 | 0.05 |
| Growth | -0.19 | 0.55 | -0.20 | 0.25 |
| Growth*Nursing | 1.27 | 0.33 | 0.36 | 0.06 |
| Growth*Carrying | 0.00 | 0.23 | 0.04 | 0.06 |
| Growth*GCs | 0.66 | 0.21 | 0.22 | 0.04 |
| Maternal ELA | -0.41 | 0.60 | -0.14 | 0.27 |

| | | | | |
|---|---|---|---|---|
| Opuntia | -0.52 | 0.54 | 0.18 | 0.24 |
| Rank | 0.34 | 0.59 | 0.34 | 0.31 |
| Parity | -0.11 | 0.92 | 0.02 | 0.41 |
| Sex | 1.46 | 0.83 | 0.25 | 0.37 |
| Infant age | -0.67 | 0.35 | -0.41 | 0.07 |
| Infant age sq | 0.61 | 0.23 | -0.06 | 0.05 |
| Group size | 0.49 | 0.59 | -0.06 | 0.27 |
| Challenges | -0.10 | 0.28 | -0.06 | 0.05 |
| Biomass | 0.33 | 0.26 | 0.24 | 0.05 |

Figure S1. Body size by infant age

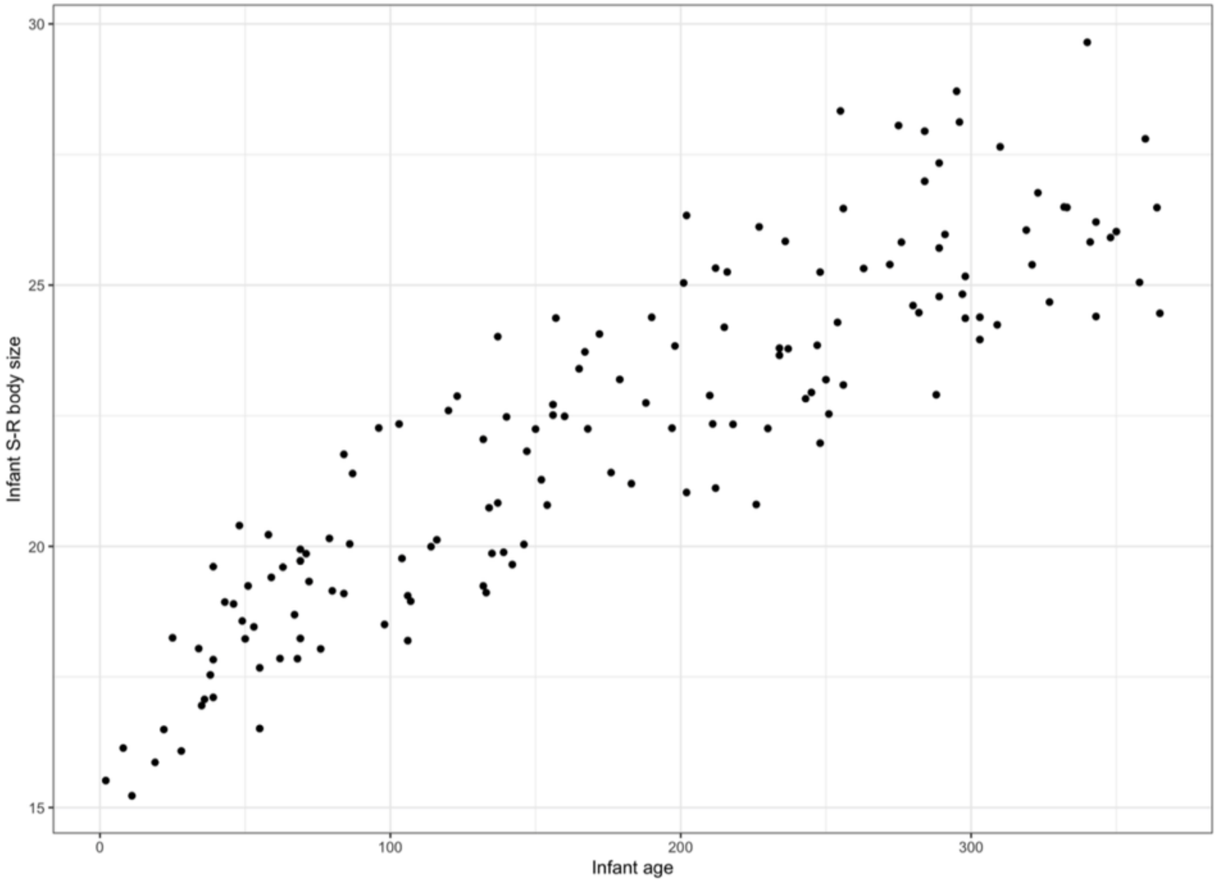


### Plots for all covariates in play, independence, and growth models:

Figure S2. The relationship between the number of infant play bouts and mother's early life adversity (ELA)

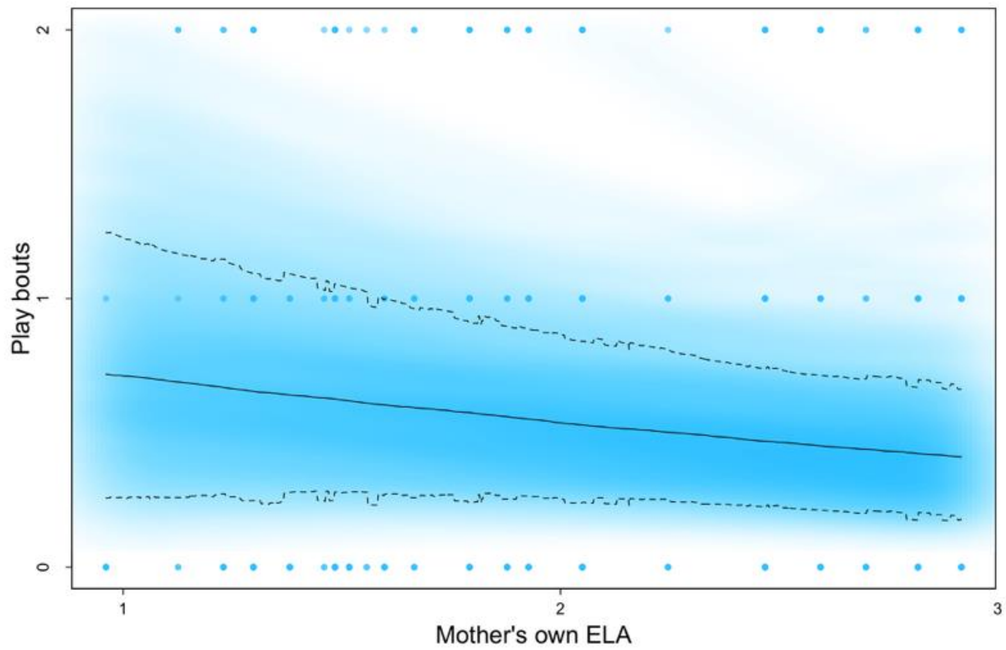


Figure S3. The relationship between the number of infant play bouts and mother's age at introduction to Opuntia stricta fruit

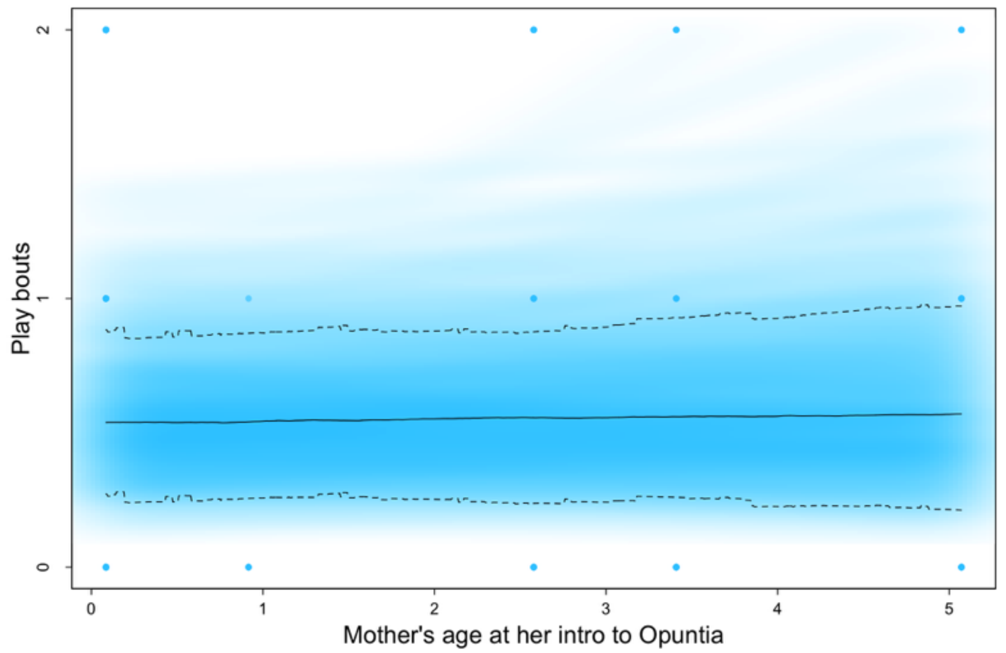


Figure S4. The relationship between the number of infant play bouts and maternal rank

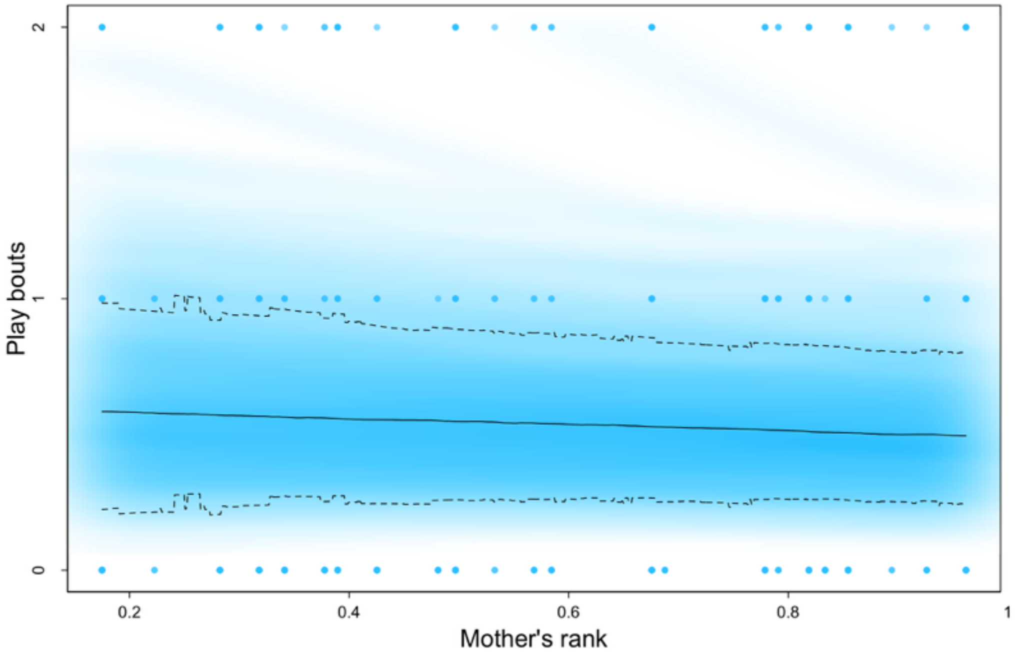


Figure S5. The relationship between the number of infant play bouts and infant age

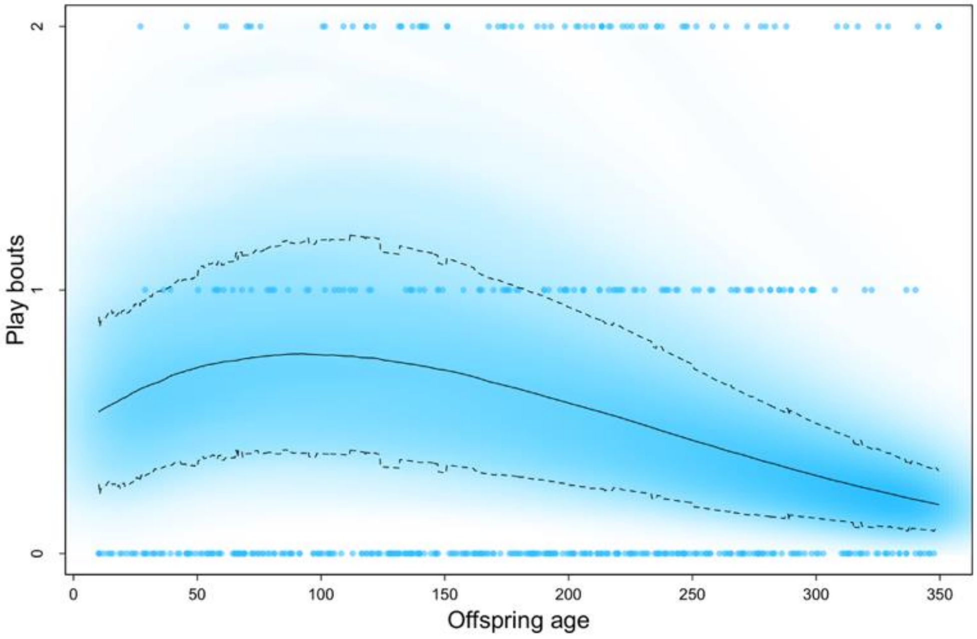


Figure S6. The relationship between the number of infant play bouts and current herbaceous biomass

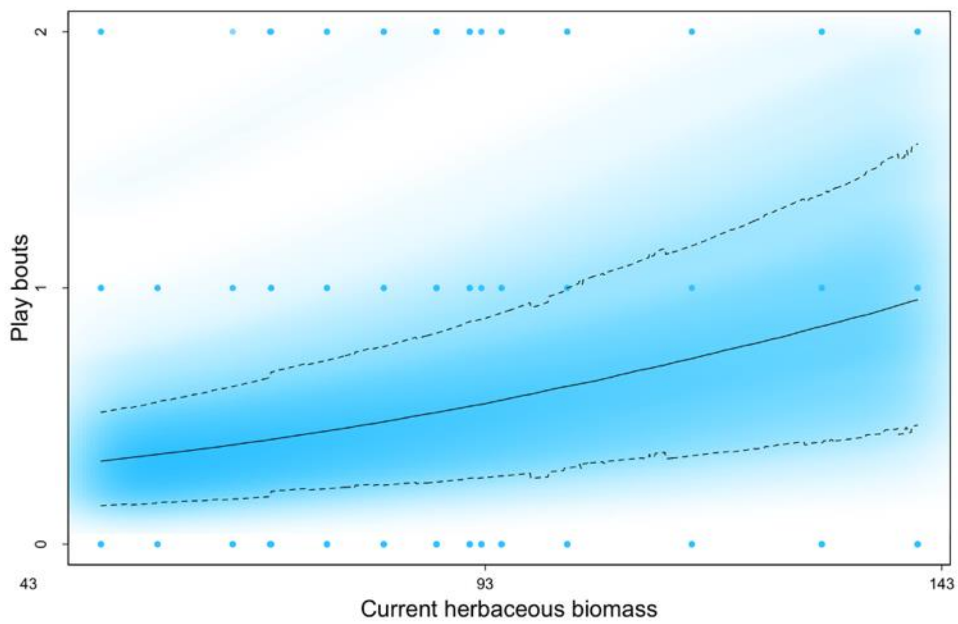


Figure S7. The relationship between the number of infant play bouts and the number of current acute environmental challenges

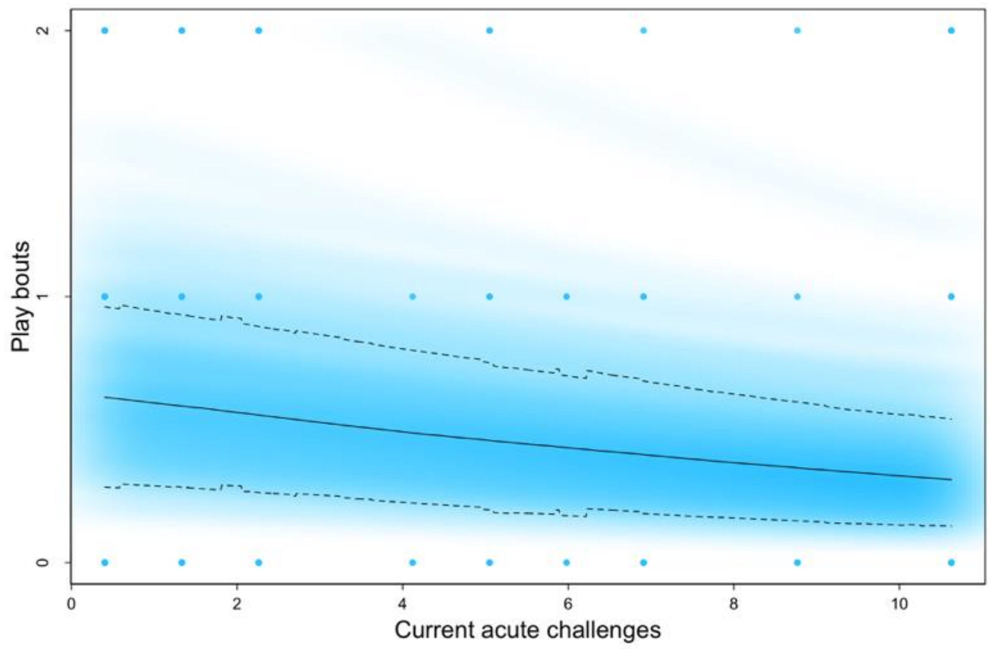


Figure S8. The relationship between the number of infant play bouts and current group size

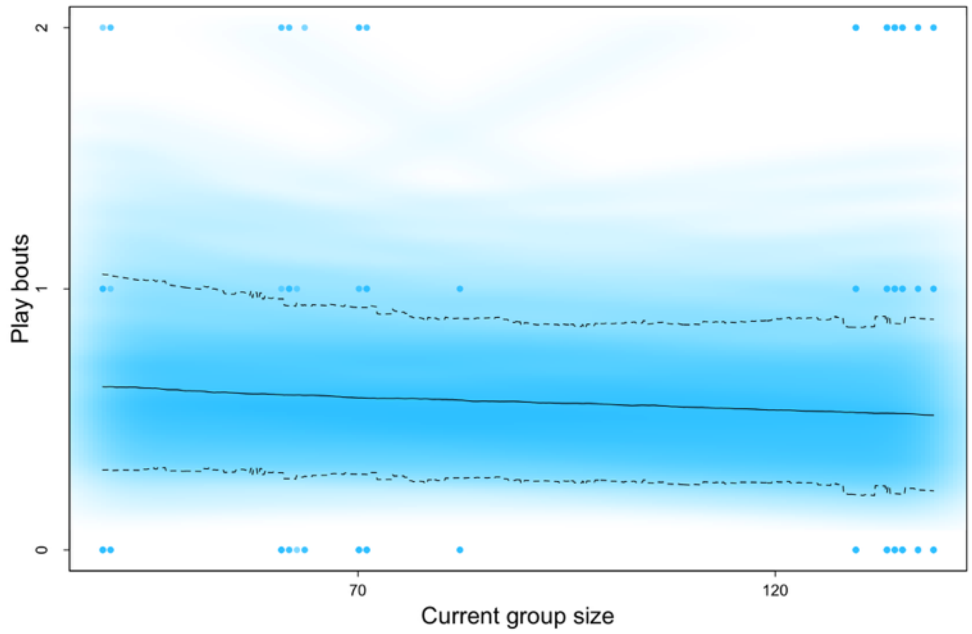


Figure S9. The relationship between the number of infant play bouts and mother's parity

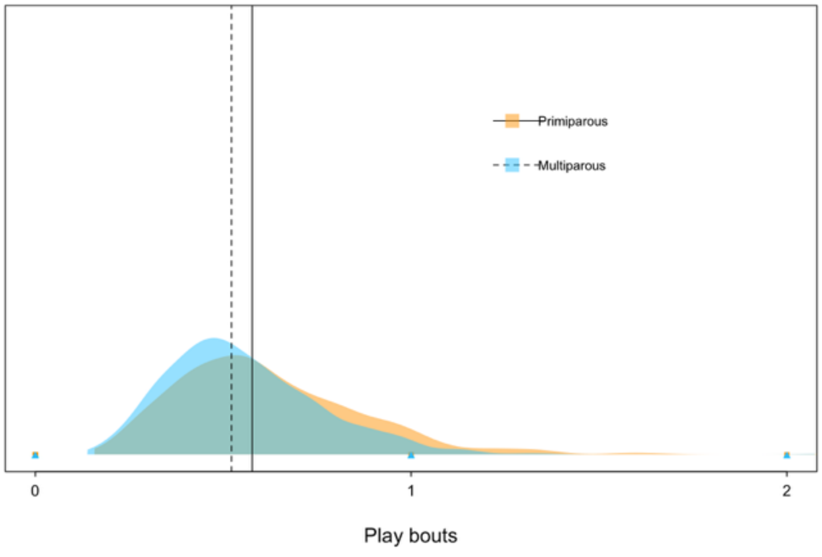


Figure S10. The relationship between the number of infant play bouts and infant sex

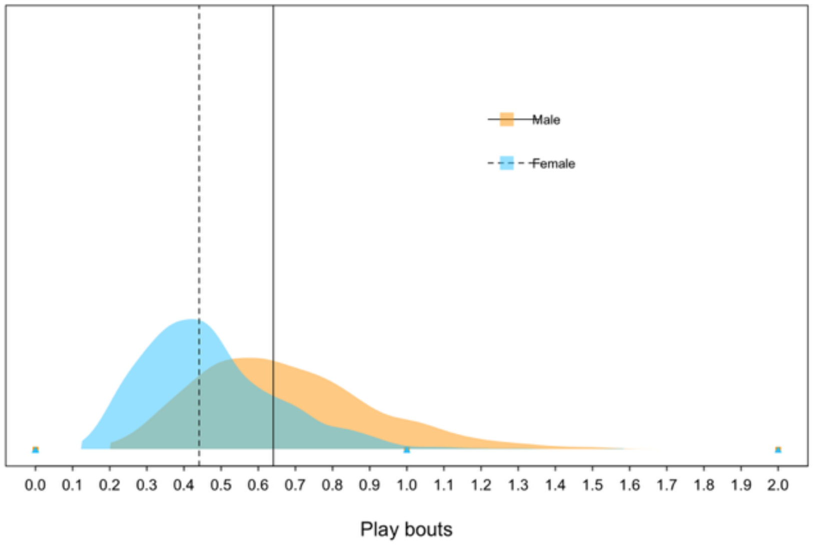


Figure S11. The relationship between the number of infant departures from mother and mother's early life adversity (ELA)

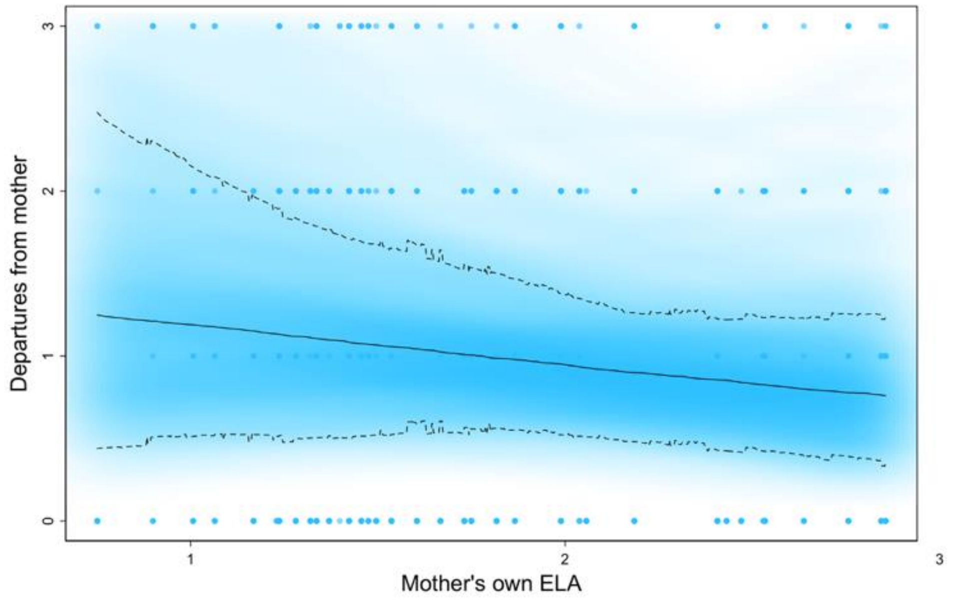


Figure S12. The relationship between the number of infant departures from mother and maternal rank

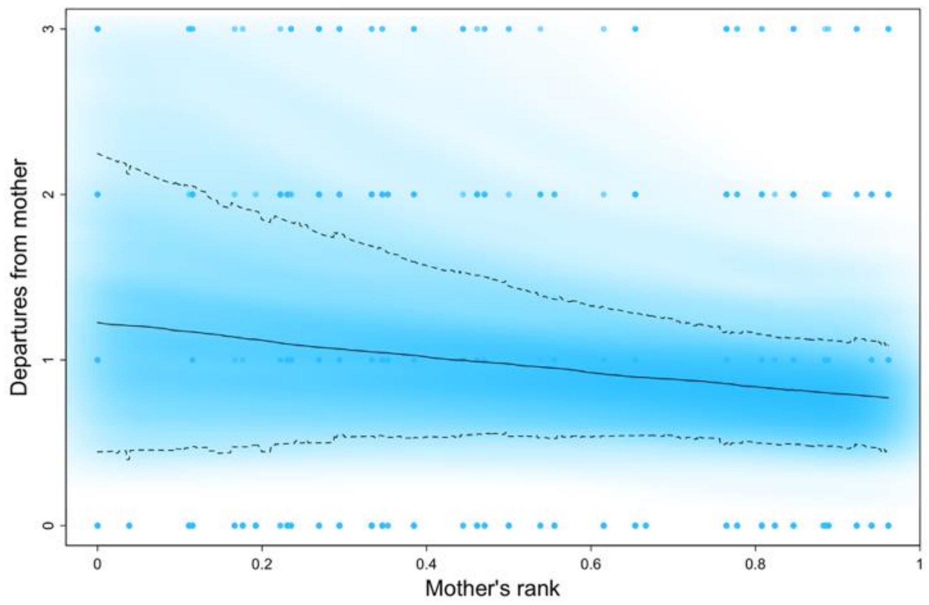


Figure S13. The relationship between the number of infant departures from mother and infant age

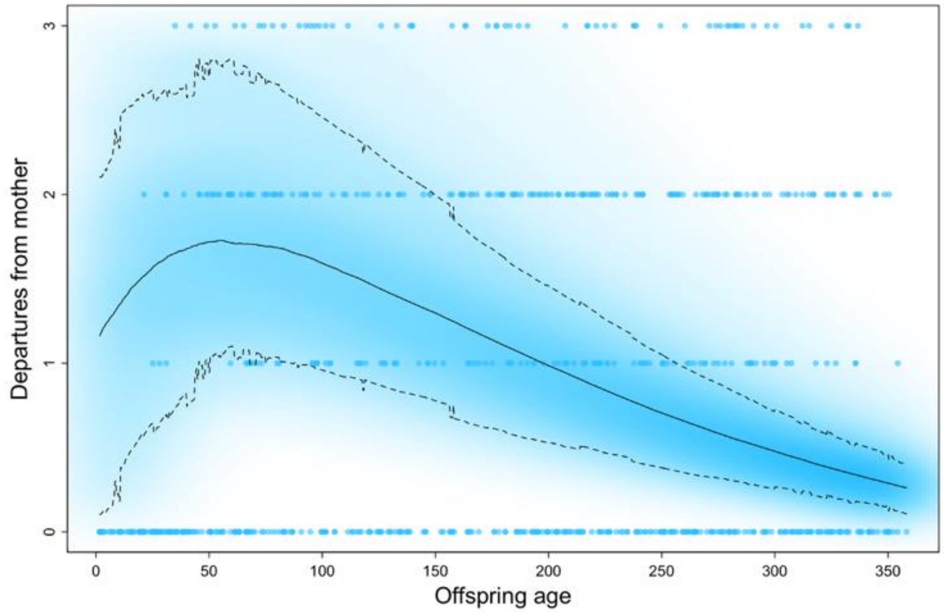


Figure S14. The relationship between the number of infant departures from mother and current herbaceous biomass

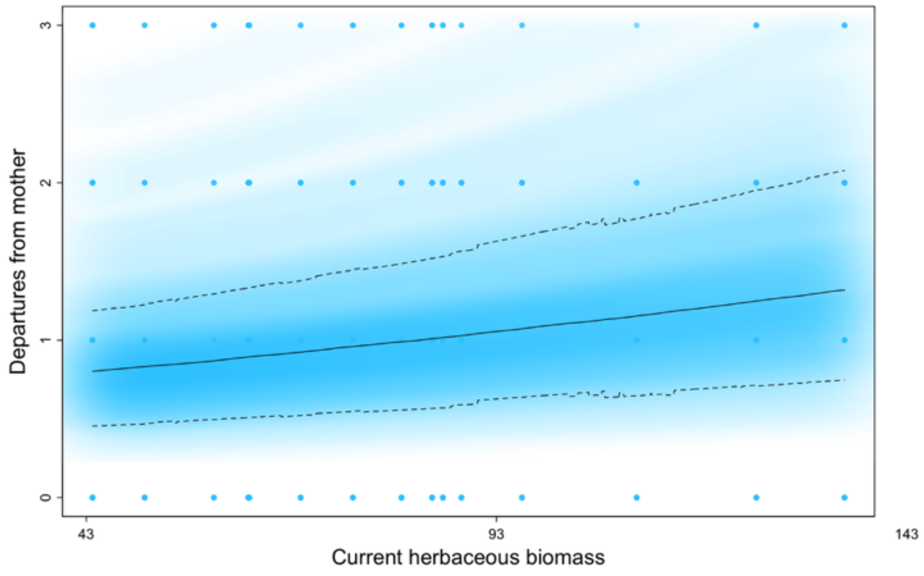


Figure S15. The relationship between the number of infant departures from mother and current acute environmental challenges

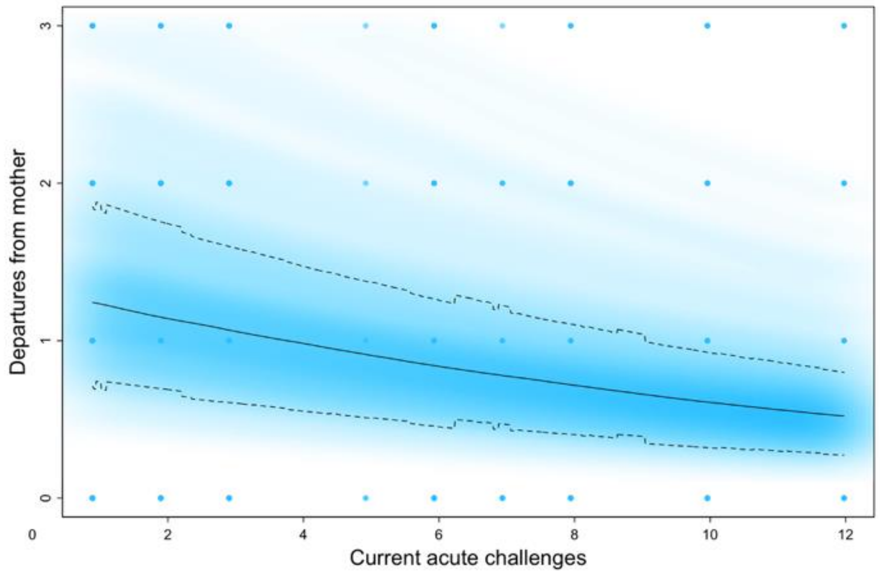


Figure S16. The relationship between the number of infant departures from mother and mother's age at introduction to Opuntia stricta fruit

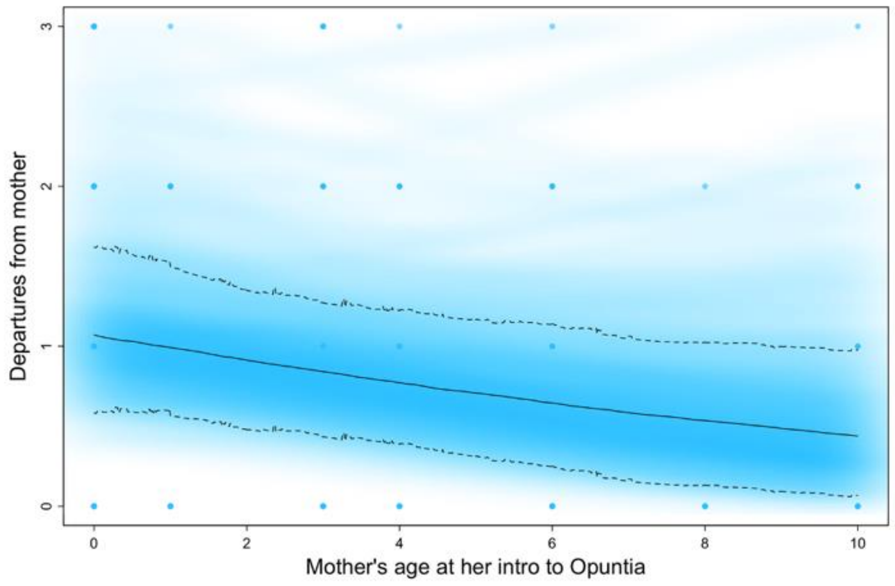


Figure S17. The relationship between the number of infant departures from mother and mother's parity

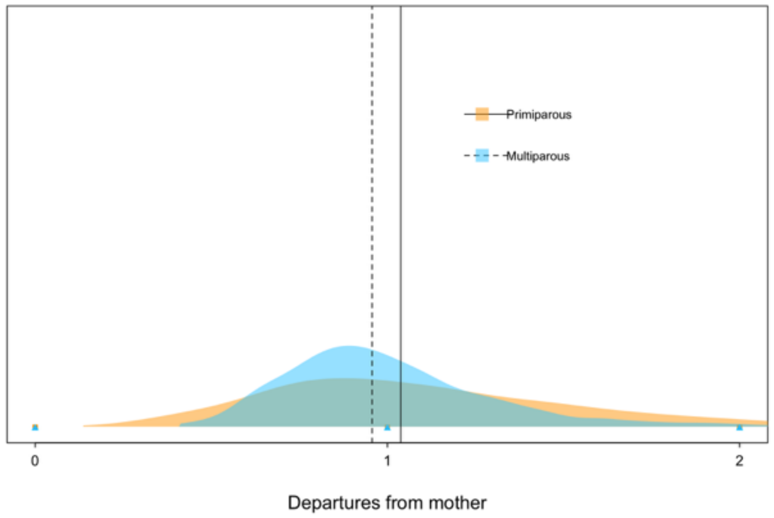


Figure S18. The relationship between the number of infant departures from mother and infant sex

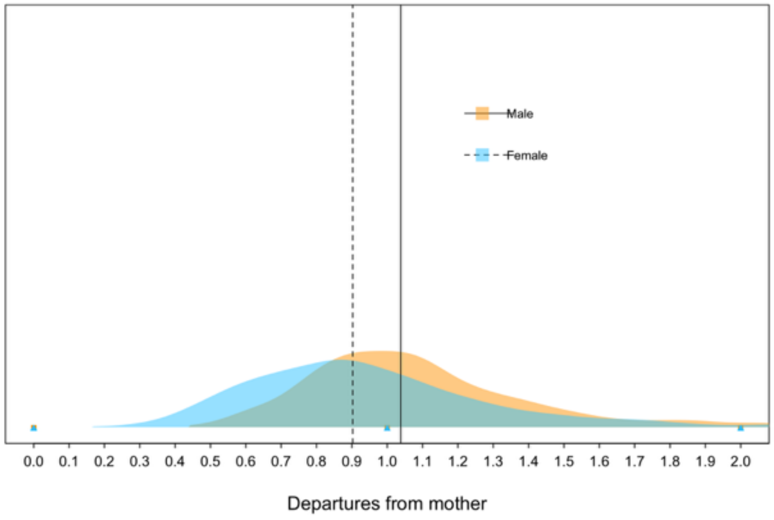


Figure S19. The relationship between infant growth and mother's early life adversity

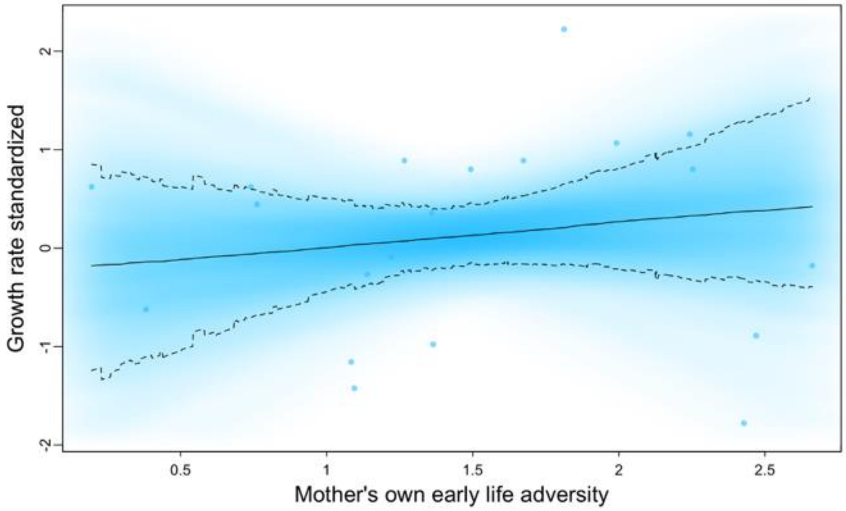


Figure S20. The relationship between infant growth and mother's introduction to Opuntia stricta fruit

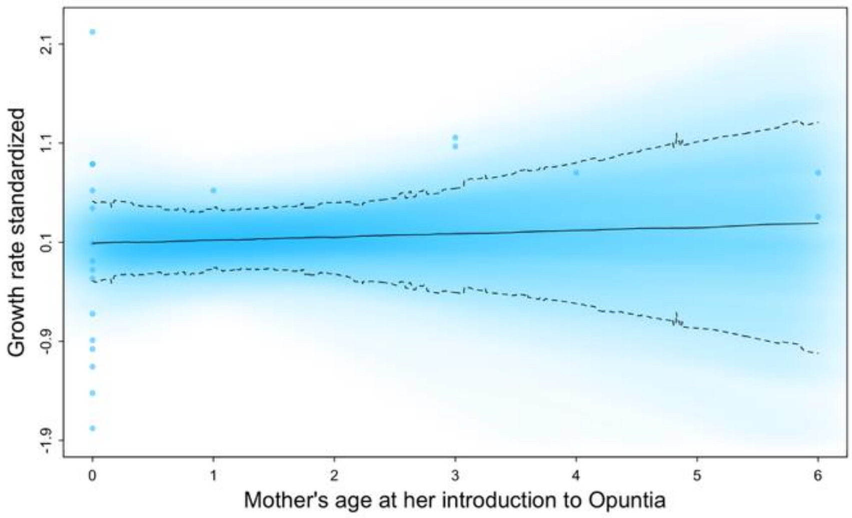


Figure S21. The relationship between infant growth and maternal rank

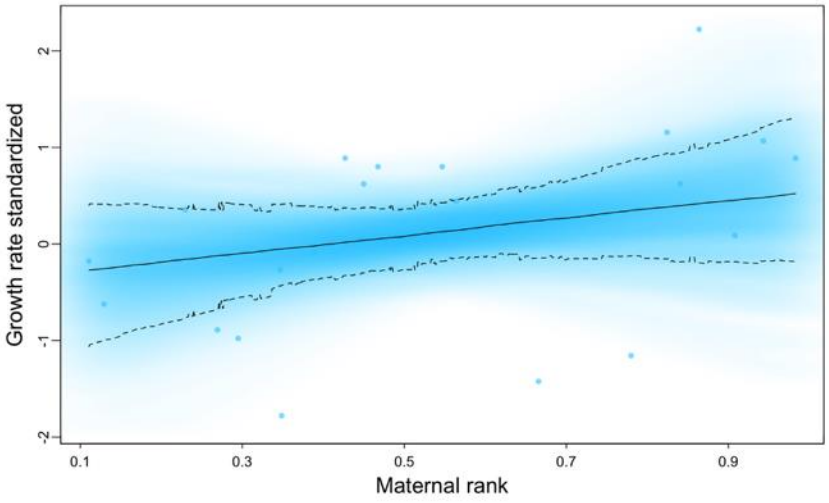


Figure S22. The relationship between infant growth and current herbaceous biomass

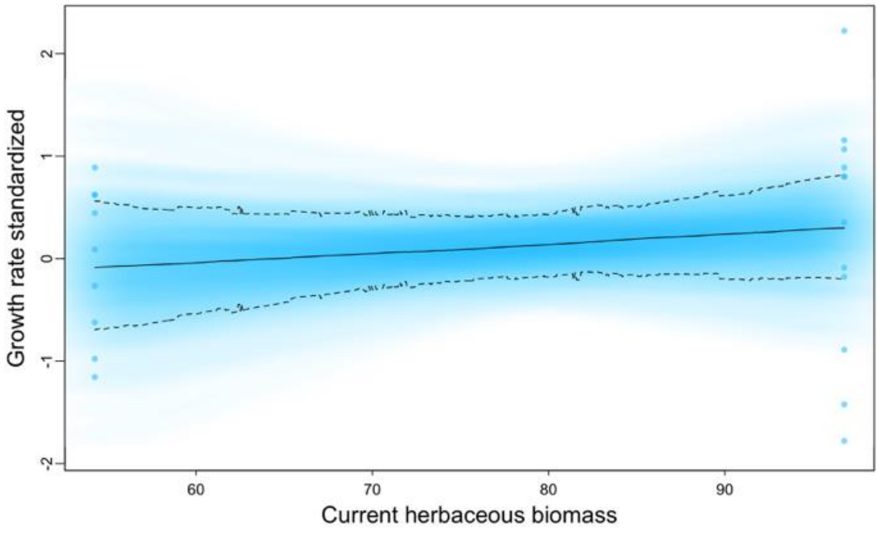


Figure S23. The relationship between infant growth and infant sex

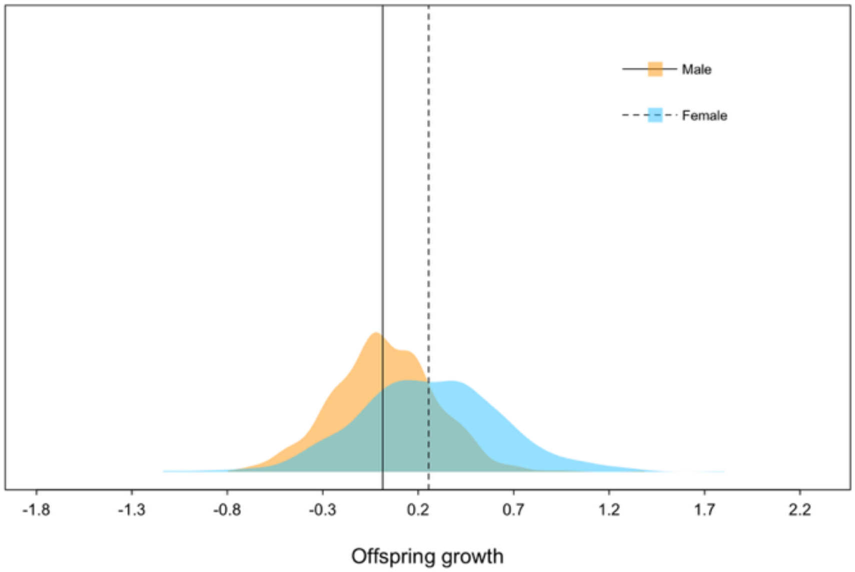


Figure S24. The relationship between infant growth and mother's parity

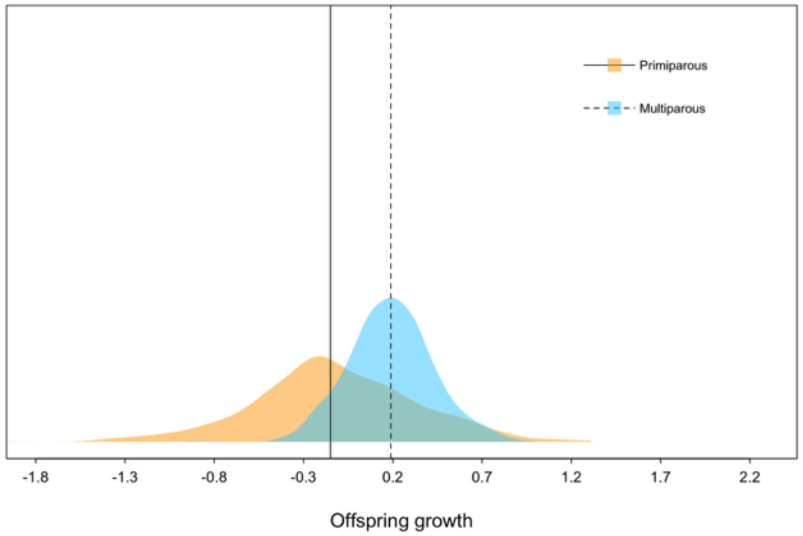

Primiparous
Multiparous
-1.8
-1.3
-0.8
-0.3
0.2
0.7
1.2
1.7
2.2
Offspring growth
